# Genomic tracking of SARS-COV-2 variants in Myanmar

**DOI:** 10.1101/2022.10.10.511623

**Authors:** Khine Zaw Oo, Zaw Win Htun, Nay Myo Aung, Ko Ko Win, Sett Paing Htoo, Thet Wai Oo, Kyaw Zawl Linn, Phyo Kyaw Aung, Myo Thiha Zaw, Kyaw Myo Tun, Kyee Myint, Ko Ko Lwin

## Abstract

**Background:** In December 2019, the COVID-19 disease started in Wuhan, China. WHO declared a pandemic on March 12, 2020, and the disease started in Myanmar on March 23, 2020. December brought variants around the world, threatening the healthcare systems. To counter those threats, Myanmar started the COVID-19 variant surveillance program in late 2020.

**Methods:** Whole genome sequencing was done six times between January 2021 and March 2022. We chose 83 samples with a PCR threshold cycle of less than 25. Then, we used MiSeq FGx for sequencing and Illumina DRAGEN COVIDSeq pipeline, command line interface, GISAID, and MEGA version 7 for data analysis.

**Result and Discussion:** January 2021 results showed no variant. The second run during the rise of cases in June 2021 showed multiple variants like Alpha, Delta, and Kappa. There is only Delta in the third run at the height of mortality in August, and Delta alone continued until the fourth run in December. After the world reported the Omicron variant in November, Myanmar started a surveillance program. The fifth run in January 2022 showed both Omicron and Delta variants. The sixth run in March 2022 showed only Omicron BA.2. Amino acid mutation at receptor binding domain (RBD) of Spike glycoprotein started since the second run coupling with high transmission, recurrence, and vaccine escape. We also found the mutation at the primer targets used in current RT-PCR platforms.

**Conclusion:** The occurrence of multiple variants and mutations claimed vigilance at ports of entry and preparedness for effective control measures. Genomic surveillance with the observation of evolutionary data is required to predict imminent threats of the current disease and diagnose emerging infectious diseases.

## Introduction

In December 2019, COVID-19 disease started in Wuhan, China (1, 2). WHO declared pandemic on March 11, 2020 (3). COVID-19 is caused by a Human Coronavirus, SARS-CoV-2. Human coronavirus (HCoV) is a member of the coronavirinae subfamily of the coronaviridae family. There were six different types of HCoV’s; two belongs to the genus Alphacoronavirus (229E, NL63) and four to Betacoronavirus (HKU1 and OC43, SARS-CoV, MERS-CoV) (3).

In Myanmar, the first wave started with two cases on March 23, 2020. There were only 374 cases and six deaths during the first wave, which ended on July 16 (4). The second wave started on August 16 in Rakhine State, followed by an outbreak in Yangon occurring among 32,351 cases with 765 fatalities. The first wave was comparatively moderate compared to regional peers, but there was a dramatic rise in the death toll during the second wave. Early June 2021 brought the third wave with a massive number of cases. Cases reached 144,157 with a death toll of 3,334, which outnumbered the capacity of quarantine centers and hospitals. At the end of the year, the total confirmed cases reached 530,834, with 19268 deaths. Omicron started on January 7, 2022, which caused a raised number of cases reaching 613,577 people with 19,434 deaths as of June 29, 2022 (4).

SARS-CoV-2 is a fast-evolving virus because of rapid and massive genetic variations. WHO classified viruses with Alpha, Beta, Gamma, Delta, and Omicron as the variant of concern (VOC) and Zeta, Eta, and Kappa as the variant of interest (VOI) (5). Since December 2020, variants have occurred worldwide, which favors the virus to be fitter for transmission. The high transmission rate caused more morbidity and mortality, endangering existing healthcare systems. All these things affected Myanmar, and she started its COVID-19 variant surveillance program in late 2020, and the Defence Services Medical Research Centre started whole genome sequencing in December 2020.

This study uses a WGS technique on the clinical samples collected during the waves above. We describe the features of the viral genome sequences from these periods, including VOCs, VOIs, and genetic variation at the spike glycoprotein coding region and primer binding sites, and draw the dynamic of viral spread.

## Materials and Methods

### Ethical Statement

The study was reviewed and approved by the institutional review board (IRB) of the Defence Services Medical Research Center (Approved ID: IRB/DSMRC/2020/15). Written informed consent form was obtained from all participants. All procedures were conducted according to the institutional guideline on responsible conduct of research.

### RNA extraction and Ct value checking

We took 159 SARS-CoV-2 PCR positive samples from clustered cases in cities and border areas during the second, third, and Omicron waves. We used QIAamp Viral RNA Mini Kit (QIAGEN, Hilden, Germany) for RNA extraction, BioFlux SARS-CoV-2 Nucleic Acid Detection Kit (Republic of China), and Applied Biosystems™ StepOnePlus™ Real-Time PCR System to check Ct values. We chose samples with a Ct value below 25 for sequencing and finally got 101 samples.

### Sequencing procedure

We performed the amplification of the SARS-CoV-2 whole genome according to the Illumina COVIDSeq RUO Assay (Illumina, USA) protocol (6). Briefly, we synthesized cDNA with the COVIDseq First Strand cDNA synthesis kit, amplified the whole viral genome selectively with Primer Pool 1 and 2, fragmented and tagged with Bead-linked transposon. Then, we added Illumina Nextera DNA CD indexes to the samples for identification before we pooled, purified, quantified, and normalized them. Finally, the MiSeq FGx sequencing system amplified them to form clusters and sequenced them using sequencing by synthesis (SBS) chemistry.

### Sequence analysis

We used Illumina DRAGEN COVIDSeq Test Pipeline to trim and assemble FASTQ files to get the FASTA format together with variant and lineage reports. The results were validated again by generating FASTA files with SC-2pipe (BioEasy Sdn Bhd, Malaysia) in the local workstation and deposited at GISAID and NCBI GenBank.

### Phylogenetic analysis

We collected 189 complete genome sequences from GISAID, which had 99.8% sequence homology with our samples. Then, we removed duplicate and incomplete sequences and finally got 162 sequences. We performed sequence alignment using Molecular Evolutionary Genetics Analysis (MEGA) software version 7 with Multiple Sequence Comparison by Log- Expectation (MUSCLE) algorithm (7), neighbor-joining analysis method, and the Kimura 2-parameter model with 1000 bootstrap replications for phylogenetic analysis.

Nucleotide and amino acid mutation sites were viewed throughout the whole viral genome length, emphasizing the Spike glycoprotein coding region to assess the effect on viral transmission and immune evasion. Mutations at ORF1ab, Nucleocapsid (N), and Envelope (E) encoding regions were also assessed, especially at primer binding sites used in PCR assays. We also collected phenotypic data, including disease severity, vaccination status, and traveling history, to correlate with viral genotypes.

## Results

We chose 83 sequences ranging in length from 28207 to 29,830 nucleotides, and each contains over 99.75% of the genome and 162 homologous sequences from 21 countries. We drew a phylogenetic tree (Fig 1), and our sequences were distributed independently from each other and were similar to strains from Europe and Asia (Fig 1-PW). Our sequences were similar to those from Malaysia during the first run (Fig 1-C and D), India, France, South Korea, Yunnan, and Romania during the second run (Fig 1-B to C), Thailand, India, South Korea, USA, and Kosovo during the third run (Fig 1-A, C, E, and F), Thailand, Malaysia, India, Austria, Germany, France, Brazil, and England during the fourth run (Fig 1-A, B, C, E, and F), Malaysia, Philippines, South Korea, India, England, and Ecuador during the fifth run(Fig 1-B, C, D, and E), and India during the sixth run (Fig 1-B, C, and D).

**Figure 1.**
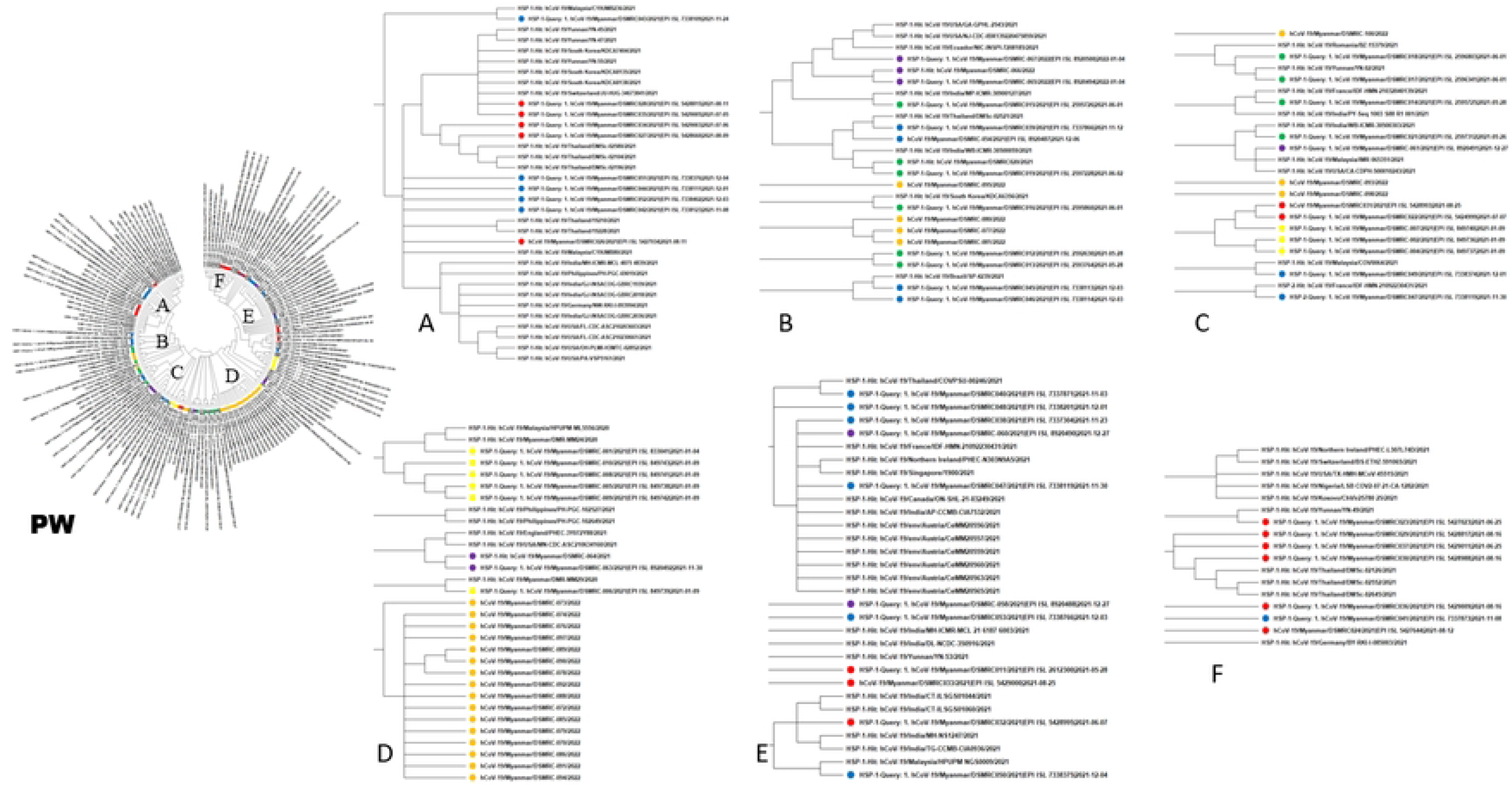
Phylogenetic tree comparing the Eighty-three Myanmar SARS-CoV-2 with 79/162 previously published sequences. Each cluster was detailed (A to F). The sequences from each run were labeled separately. (Yellow = First Run, Green = Second Run, Red = Third Run, Blue = Fourth Run, Purple = Fifth Run and Orange = Sixth Run)

All our samples showed D614G mutation since the first run with additional Q677H mutation at 5 out of 9 samples. No variant of concern (VOC) or variant of interest (VOI) existed (Table 1). After six months, we did the second run and found 7 VOCs (2 Alpha and 5 Delta) and 4 VOIs (Kappa). Alpha variants showed N501Y mutation at RBD, while the Delta showed L452R and T478K mutations at RBD with P681H mutation at the furin binding site (FBS). Like Delta variants, Kappa showed L452R and P681H mutations, but there was E484Q instead of T478K mutation (Table 2). The third and fourth runs got 15 and 16 complete sequences, showing only the Delta variant with the same mutation as before. Four Delta remained present during the fifth run, together with 5 Omicron variants. The sixth run in March 2022, with 23 samples, showed only Omicron (Table 3), all having multiple mutations at RBD and FBS.

**Table 1.** PANGO Lineage, Clade, Variant, Spike protein mutations of First and Second Run (During the Second Wave)

| Sample Name | PANGO Lineage | Nextstrain Clade | Spike Mutation from RBD to FBS | Similar Countries |
| --- | --- | --- | --- | --- |
| DSMRC-001 | B.1.36.16 | 20A | D614G | Malaysia |
| DSMRC-002 | B.1.36.16 | 20A | D614G | Malaysia |
| DSMRC-004 | B.1.36.16 | 20A | D614G | Malaysia |
| DSMRC-005 | B.1.36.16 | 20A | D614G, Q677H | Malaysia |
| DSMRC-006 | B.1.36.16 | 20A | D614G | Malaysia |
| DSMRC-007 | B.1.36.16 | 20A | D614G, Q677H | Malaysia |
| DSMRC-008 | B.1.36.16 | 20A | D614G, Q677H | Malaysia |
| DSMRC-009 | B.1.36.16 | 20A | D614G, Q677H | Malaysia |
| DSMRC-010 | B.1.36.16 | 20A | D614G, Q677H | Malaysia |
| DSMRC-011 | B.1.617.2 | 21I (Delta) | D614G, L452R | France |
| DSMRC-012 | B.1.1.7 | 20I (Alpha, V1) | D614G, N501Y | India |
| DSMRC-013 | B.1.617.1 | 21B (Kappa) | D614G, L452R, E484Q | India |
| DSMRC-014 | B.1.1.7 | 20I (Alpha, V1) | D614G, N501Y | France |
| DSMRC-015 | B.1.617.2 | 21A (Delta) | D614G, L452R, T478K, P681H | India |
| DSMRC-016 | B.1.617.2 | 21A (Delta) | D614G, L452R, T478K, P681H | South Korea |
| DSMRC-017 | B.1.617.2 | 21A (Delta) | D614G, L452R, T478K, P681H | Romania |
| DSMRC-018 | B.1.617.2 | 21A (Delta) | D614G, L452R, T478K, P681H | Yunnan |
| DSMRC-019 | B.1.617.1 | 21B (Kappa) | D614G, L452R, E484Q, P681H | India |
| DSMRC-020 | B.1.617.1 | 21B (Kappa) | D614G, L452R, E484Q, P681H | India |
| DSMRC-021 | B.1.617.1 | 21B (Kappa) | D614G, L452R, E484Q,<br>P681H | India |

**Table 2.** PANGO Lineage, Clade, Variant, Spike protein mutations of Third, Fourth, and Fifth Run (During the Third Wave)

| Sample Name | PANGO Lineage | Nextstrain Clade | Spike Mutation from RBD to FBS | Similar Countries |
| --- | --- | --- | --- | --- |
| DSMRC-022 | AY.30 | 21A (Delta) | D614G, L452R, T478K, P681H | Thailand |
| DSMRC-023 | B.1.617.2 | 21A (Delta) | D614G, L452R, T478K, P681H | South Korea |
| DSMRC-024 | B.1.617.2 | 21I (Delta) | D614G, L452R, T478K, P681H | India |
| DSMRC-026 | B.1.617.2 | 21I (Delta) | D614G, L452R, T478K, P681H | India |
| DSMRC-027 | AY.30 | 21A (Delta) | D614G, L452R, T478K, P681H | Thailand |
| DSMRC-028 | B.1.617.2 | 21A (Delta) | D614G, L452R, T478K, P681H | South Korea |
| DSMRC-029 | AY.30 | 21A (Delta) | D614G, L452R, T478K, P681H | Thailand |
| DSMRC-030 | AY.30 | 21A (Delta) | D614G, L452R, T478K, P681H | India |
| DSMRC-031 | AY.30 | 21A (Delta) | D614G, L452R, T478K, P681H | Thailand |
| DSMRC-032 | B.1.617.2 | 21A (Delta) | D614G, L452R, T478K, P681H | India |
| DSMRC-033 | B.1.617.2 | 21A (Delta) | D614G, L452R, T478K, P681H | USA |
| DSMRC-034 | AY.30 | 21A (Delta) | D614G, L452R, T478K, P681H | Thailand |
| DSMRC-035 | B.1.617.2 | 21A (Delta) | D614G, L452R, T478K, P681H | South Korea |
| DSMRC-036 | B.1.617.2 | 21A (Delta) | D614G, L452R, T478K, P681H | Kosovo |
| DSMRC-037 | AY.30 | 21A (Delta) | D614G, L452R, T478K, Q675H, P681H | Thailand |
| DSMRC-038 | B.1.617.2 | 21J (Delta) | D614G, L452R, T478K, P681H | Austria |
| DSMRC-039 | AY.85 | 21J (Delta) | D614G, L452R, T478K, P681H | Thailand |
| DSMRC-040 | AY.85 | 21J (Delta) | D614G, L452R, T478K, P681H | Thailand |
| DSMRC-041 | AY.60 | 21I (Delta) | D614G, L452R, T478K, P681H | Germany |
| DSMRC-042 | AY.60 | 21I (Delta) | D614G, L452R, T478K, P681H | Germany |
| DSMRC-043 | AY.79 | 21J (Delta) | D614G, L452R, T478K, P681H | Malaysia |
| DSMRC-044 | AY.60 | 21I (Delta) | D614G, L452R, T478K, P681H | Germany |
| DSMRC-045 | B.1.617.2 | 21J (Delta) | D614G, L452R, T478K, P681H | Brazil |

**Table 2. PANGO Lineage, Clade, Variant, Spike protein mutations of Third, Fourth,** 143 **and Fifth Run (During the Third Wave) (cont.)**
| Sample Name | PANGO Lineage | Nextstrain Clade | Spike Mutation from RBD to FBS | Similar Countries |
| --- | --- | --- | --- | --- |
| DSMRC-046 | B.1.617.2 | 21J (Delta) | D614G, L452R, T478K, P681H | Brazil |
| DSMRC-047 | AY.95 | 21J (Delta) | D614G, L452R, T478K, P681H | France |
| DSMRC-048 | B.1.617.2 | 21J (Delta) | D614G, L452R, T478K, P681H | Canada |
| DSMRC-049 | B.1.617.2 | 21J (Delta) | D614G, L452R, T478K, P681H | Malaysia |
| DSMRC-050 | AY.59 | 21I (Delta) | D614G, L452R, T478K, P681H | Malaysia |
| DSMRC-051 | AY.59 | 21I (Delta) | D614G, L452R, T478K, P681H | Malaysia |
| DSMRC-052 | AY.60 | 21I (Delta) | D614G, L452R, T478K, P681H | India |
| DSMRC-053 | AY.60 | 21I (Delta) | D614G, L452R, T478K, Q414R, P681H | England |
| DSMRC-054 | B.1.617.2 | 21J (Delta) | D614G, L452R, T478K, P681H | South Korea |
| DSMRC-058 | AY.60 | 21J (Delta) | D614G, L452R, T478K, P681H | India |
| DSMRC-060 | AY.79 | 21J (Delta) | D614G, L452R, T478K, P681H | Malaysia |
| DSMRC-061 | AY.79 | 21J (Delta) | D614G, L452R, T478K, P681H | Malaysia |
| DSMRC-063 | BA.1 | 21K (Omicron) | D614G, D796Y, E484A, G339D, G446S, G496S, K417N, N440K, N501Y, N764K, P681H, Q493R, Q498R, S371L, S373P, S375F, S477N, T478K, Y505H | England |
| DSMRC-064 | BA.1 | 21K (Omicron) | D614G, D796Y, E484A, G339D, G446S, G496S, K417N, N440K, N501Y, N764K, P681H, Q493R, Q498R, S371L, S373P, S375F, S477N, T478K, Y505H | England |
| DSMRC-065 | BA.2 | 21L (Omicron) | D614G, D796Y, E484A, G339D, G496S, K417N, R408S, N501Y, N764K, P681H, Q493R, Q498R, S371L, S373P, S375F, S477N, T478K, Y505H | Philippine |

**Table 2. PANGO Lineage, Clade, Variant, Spike protein mutations of Third, Fourth, and Fifth Run (During the Third Wave) (cont.)**
| Sample Name | PANGO Lineage | Nextstrain Clade | Spike Mutation from RBD to FBS | Similar Countries |
| --- | --- | --- | --- | --- |
| DSMRC-066 | BA.2 | 21L (Omicron) | D614G, D796Y, E484A, G339D, G496S, K417N, R408S, N501Y, N764K, P681H, Q493R, Q498R, S371L, S373P, S375F, S477N, T478K, Y505H | Philippine |
| DSMRC-067 | BA.1 | 21K (Omicron) | D614G, D796Y, E484A, G339D, R346K, G446S, G496S, K417N, N440K, N501Y, N764K, P681H, Q493R, Q498R, S371L, S373P, S375F, S477N, T478K, Y505H | Ecuador |

**Table 3.** PANGO Lineage, Clade, Variant, Spike protein mutations of Sixth Run (During the last wave)

| Sample Name | PANGO Lineage | Next strain Clade | Spike Mutation from RBD to FBS | Similar Countries |
| --- | --- | --- | --- | --- |
| DSMRC070 | BA.2 | 21L (Omicron) | D405N, D614G, D796Y, E484A, G339D, H655Y, K417N, N440K, N501Y, N679K, N764K, P681H, Q493R, Q498R, R408S, S371F, S373P, S375F, S477N, T376A, T478K, Y505H |  |
| DSMRC-072 | BA.2 | 21L (Omicron) | D405N, D614G, D796Y, E484A, G339D, H655Y, K417N, N440K, N501Y, N679K, N764K, P681H, Q493R, Q498R, R408S, S371F, S373P, S375F, S477N, T376A, T478K, Y505H |  |
| DSMRC-073 | BA.2 | 21L (Omicron) | D405N, D614G, D796Y, E484A, G339D, H655Y, K417N, N440K, N501Y, N679K, N764K, P681H, Q493R, Q498R, R408S, S371F, S373P, S375F, S477N, T376A, T478K, Y505H |  |
| DSMRC-074 | BA.2 | 21L (Omicron) | D405N, D614G, D796Y, E484A, G339D, H655Y, K417N, N440K, N501Y, N679K, N764K, P681H, Q493R, Q498R, R408S, S371F, S373P, S375F, S477N, T376A, T478K, Y505H |  |
| DSMRC-076 | BA.2 | 21L (Omicron) | D614G, D796Y, E484A, G339D, H655Y, K417N, N440K, N501Y, N679K, N764K, P681H, Q493R, Q498R, R408S, S371F, S373P, S375F, S477N, T376A, T478K, Y505H |  |
| DSMRC-077 | BA.2 | 21L (Omicron) | D614G, D796Y, E484A, H655Y, N501Y, N679K, N764K, P681H, Q493R, Q498R, R408S, S477N, T478K, Y505H |  |

**Table 3. PANGO Lineage, Clade, Variant, Spike protein mutations of Sixth Run** 153 **(During the last wave) (cont.)**
| Sample Name | PANGO Lineage | Next strain Clade | Spike Mutation from RBD to FBS | Similar Countries |
| --- | --- | --- | --- | --- |
| DSMRC-078 | BA.2 | 21L (Omicron) | D405N, D614G, D796Y, E484A, G339D, H655Y, K417N, N440K, N501Y, N679K, N764K, P681H, Q493R, Q498R, R408S, S371F, S373P, S375F, S477N, T376A, T478K, Y505H |  |
| DSMRC-079 | BA.2 | 21L (Omicron) | D405N, D614G, D796Y, E484A, G339D, H655Y, K417N, N440K, N501Y, N679K, N764K, P681H, Q493R, Q498R, R408S, S371F, S373P, S375F, S477N, T376A, T478K, Y505H |  |
| DSMRC-080 | BA.2 | 21L (Omicron) | D405N, D614G, D796Y, E484A, G339D, H655Y, K417N, N440K, N501Y, N679K, N764K, P681H, Q493R, Q498R, R408S, S371F, S373P, S375F, S477N, T376A, T478K, Y505H |  |
| DSMRC-081 | BA.2 | 21L (Omicron) | D614G, D796Y, E484A, G339D, H655Y, K417N, N440K, N501Y, N679K, N764K, P681H, Q493R, Q498R, R408S, S371F, S373P, S375F, S477N, T376A, T478K, Y505H |  |
| DSMRC-085 | BA.2 | 21L (Omicron) | D405N, D614G, D796Y, E484A, G339D, H655Y, K417N, N440K, N501Y, N679K, N764K, P681H, Q493R, Q498R, R408S, S371F, S373P, S375F, S477N, T376A, T478K, Y505H |  |
| DSMRC-086 | BA.2 | 21L (Omicron) | D405N, D614G, D796Y, E484A, G339D, H655Y, K417N, N440K, N501Y, N679K, N764K, P681H, Q493R, Q498R, R408S, S371F, S373P, S375F, S477N, T376A, T478K, Y505H |  |
| DSMRC-088 | BA.2 | 21L (Omicron) | D614G, D796Y, E484A, G339D, H655Y, K417N, N440K, N501Y, N679K, N764K, P681H, Q493R, Q498R, R408S, S371F, S373P, S375F, S477N, T376A, T478K, Y505H |  |
| DSMRC-089 | BA.2 | 21L (Omicron) | D405N, D614G, D796Y, E484A, G339D, H655Y, K417N, N440K, N501Y, N679K, N764K, P681H, Q493R, Q498R, R408S, S371F, S373P, S375F, S477N, T376A, T478K, Y505H |  |
| DSMRC-090 | BA.2 | 21L (Omicron) | D405N, D614G, D796Y, E484A, G339D, H655Y, K417N, N440K, N501Y, N679K, N764K, P681H, Q493R, Q498R, R408S, S371F, S373P, S375F, S477N, T376A, T478K, Y505H |  |

**Table 3. PANGO Lineage, Clade, Variant, Spike protein mutations of Sixth Run** **(During the last wave) (cont.)**
| Sample Name | PANGO Lineage | Next strain Clade | Spike Mutation from RBD to FBS | Similar Countries |
| --- | --- | --- | --- | --- |
| DSMRC-091 | BA.2 | 21L (Omicron) | D405N, D614G, D796Y, E484A, G339D, H655Y, K417N, N440K, N501Y, N679K, N764K, P681H, Q493R, Q498R, R408S, S371F, S373P, S375F, S477N, T376A, T478K, Y505H |  |
| DSMRC-092 | BA.2 | 21L (Omicron) | D405N, D614G, D796Y, E484A, G339D, H655Y, K417N, N440K, N501Y, N679K, N764K, P681H, Q493R, Q498R, R408S, S371F, S373P, S375F, S477N, T376A, T478K, Y505H |  |
| DSMRC-093 | BA.2 | 21L (Omicron) | D405N, D614G, D796Y, E484A, G339D, H655Y, K417N, N440K, N501Y, N679K, N764K, P681H, Q493R, Q498R, R408S, S371F, S373P, S375F, S477N, T376A, T478K, Y505H |  |
| DSMRC-094 | BA.2 | 21L (Omicron) | D405N, D614G, D796Y, E484A, G339D, H655Y, K417N, N440K, N501Y, N679K, N764K, P681H, Q493R, Q498R, R408S, S371F, S373P, S375F, S477N, T376A, T478K, Y505H |  |
| DSMRC-095 | BA.2 | 21L (Omicron) | D405N, D614G, D796Y, E484A, G339D, H655Y, K417N, N440K, N501Y, N679K, N764K, P681H, Q493R, Q498R, R408S, S371F, S373P, S375F, S477N, T376A, T478K, Y505H |  |
| DSMRC-097 | BA.2 | 21L (Omicron) | D405N, D614G, D796Y, E484A, G339D, H655Y, K417N, N440K, N501Y, N679K, N764K, P681H, Q493R, Q498R, R408S, S371F, S373P, S375F, S477N, S704L, T376A, T478K, Y505H |  |
| DSMRC-098 | BA.2 | 21L (Omicron) | D614G, D796Y, E484A, H655Y, N501Y, N679K, P681H, Q493R, Q498R, S477N, T478K, Y505H |  |
| DSMRC-100 | BA.2 | 21L (Omicron) | D614G, D796Y, E484A, H655Y, N501Y, N679K, P681H, Q493R, Q498R, S477N, T478K, Y505H |  |

Three out of four Delta variants during the fifth run (DSMRC 054, 058, 060) showed Nucleocapsid G28881T, and DSMRC 058, 060 showed ORF1ab G15451A nucleotide substitutions. Three Omicron variants (DSMRC 063, 064, 067) showed trinucleotide mutations (G28881A, G28882A, G28883C) at the Nucleocapsid coding region and Envelope C26270T mutation.

All our sequences belonged to non-variant and four variants, and 15 strains. Generally, they were distributed equally with each other and worldwide sequences. However, the third and fourth runs with the Delta variant only became more homologous with them.

## Discussion

Many variants were found in this study and were like India, England, and worldwide. We found similar sequences from Asia and worldwide without anyone from Africa and South America. Since late 2021, the Delta variant dominated until the appearance of Omicron in January 2022. Delta variant took over other global variances after its emergence in India in December 2020 (8).

Phylogenetic analysis showed that our sequences were distributed independently and clustered together with those from 21 countries. These results showed complex geographical and temporal transmission patterns and a high virus mutation rate. Our sequences became more homologous with worldwide sequences during the third and fourth runs. Clustered with those from Europe (France, England) showed rapid disease transmission in today’s world. Clustered with other Asian countries (India, Malaysia, and Thailand) showed the border crossing of disease through land and air. Sequences had several nucleotide changes compared to the first published COVID-19 sequence from Wuhan, China (NC045512.2) (9).

D614G mutation, found in January 2021, was first detected in January 2020 in China and Germany. It outnumbered the initial coronavirus by April/May (10). G614 variant had improved receptor binding and transmission, becoming the dominant global strain (9). Molecular courting also indicated that this novel mutation ascended early in the pandemic. Genetic drift caused a high rate of a specific variant without selective advantage (11). Q677H mutation found in some of our samples might affect the stability of the spike protein as it lay near the furin cleavage site, but there was no evidence of its effect on the pathogenicity (12). There was no difference in severity between patients with Q677H and without.

N501Y mutation seen in Alpha, and L452R and T478K in Delta variant, lied at RBD. In the UK, South Africa, and Brazil, the N501Y mutation had increased binding affinity to ACE2. L452R showed reduced neutralization by several monoclonal antibodies (mAbs) (13–15) and convalescent plasma (13). L452R independently appeared in different lineages between December 2020 and February 2021. This substitution might result from viral adaptation due to increased immunity in the population (9). T478K mutation at the interface complex with human ACE2 might also affect the affinity with human cells and viral infectivity (16).

Kristian Andersen identified a notable feature: polybasic, furin, binding at position 681-688, and subsequent cleavage of Spike protein worsened its stability. Cleavage of subunits increased binding affinity to ACE2 receptor markedly (17). Unfortunately, all Delta and Kappa variants developed with P681H mutation at FBS and would cause more transmission.

Kappa variants also showed E484Q mutation at the RBD’s receptor-binding motif (RBM). As the central functional motif, RBM directly binds the human ACE2 receptor (18). E484N and E484K escape from several mAb and antibodies in convalescent plasma (15, 19). Due to the presence of both L452R and E484Q mutations, our Kappa variant would affect the antigenicity and subsequent immune protection (18, 19). Investigations using pseudoviruses with RBD mutations revealed that the neutralizing activity of plasma from vaccinated people decreased onefold to thrice against N501Y and E484K (20).

Finally, Omicron came up with multiple mutations at RBD and FBS, which would cause the virus to be more transmissible and affect the host’s immune protection. Clinical data showed rapid transmission without evidence of disease severity and a high mortality rate. K417 is the epitope for class 1 and class 2 antibodies (21), but mutation would affect only class 1 antibody binding. So, the K417 mutation found in this study would have less effect on the polyclonal antibody response, responsible for class 2 antibody binding. They were more vulnerable to substitutions like E484K (22). K417N and K417T had lower ACE2-binding affinity in addition to their antigenic impact (23).

Currently, we are using Daan Gene and Bioflux RT-PCR and cobas6800 platforms to diagnose COVID-19. Some Delta variants showed nucleotide substitution at China CDC forward primer binding site at Nucleocapsid coding region (28881-28902) but not at the ORF1ab region. They also showed substitution at Charite forward primer binding site at ORF1b (15431-15452) but not at the Envelope coding region. Some Omicron variants showed substitution at China CDC forward primer binding site at the Nucleocapsid region (28881-28902) but not at the ORF1ab region. They also showed substitution at Charite forward primer binding site at the Envelope region (26269-26294) but not at the ORF1b region. These show that our molecular platforms are still effective for diagnosing COVID-19 (24).

Distribution of our sequences across the whole phylogenetic tree constructed with sequences from 21 countries worldwide, inferring globalization and high transmissibility of disease, warned us to keep stricter vigilance of imported cases. Sequence homology with Europe and Malaysia indicated keeping track of disease transmission patterns and ports of entry from these regions. Ongoing transmission from India from the First to the Sixth Run pointed out not to emphasize only the eastern border but to exercise more containment strategy at the west.

Multiple variants inland claimed sustainable plan and preparation for diagnosis, treatment, prevention, and control measures against repeatedly emerging diseases. Genomic data during waves of world outbreaks showed the dominance of the fittest Delta strain over other strains. This domination predicted the inevitable importation of this strain, and we should use this information effectively to prepare for future control measures. Detailed information about mutations at receptor binding sites, immune epitopes, and primer binding sites offered important intelligence data to combat the novel virus. We should maintain our efforts in genomic surveillance. Finally, the whole genome sequencing platform in this study opens our capability for diagnosing emerging infectious diseases in the upcoming days in the challenging world.

## Conclusion

Multiple variants and mutations have necessitated increased vigilance at ports of entry and the development of effective control measures. Genomic surveillance with the observation of evolutionary data is required to predict imminent threats of the current disease and diagnose emerging infectious diseases.

## Notes

### Competing Interest Statement

The authors have declared no competing interest.

